# Vitronectin acts as a key regulator of adhesion and migration in human umbilical cord-derived MSCs under different stress conditions

**DOI:** 10.1101/2021.11.18.468035

**Authors:** Ankita Sen, Malancha Ta

## Abstract

Mesenchymal stem cell (MSC)-based therapy gets compromised as adverse microenvironmental conditions like nutrient deprivation, ischemia, hypoxia at the target site affect migration, engraftment and viability, of MSCs post transplantation. To improve the treatment efficacy, it is critical to identify factors involved in regulating migration and adhesion of MSCs under such microenvironmental stress conditions. In our study, we observed that human Wharton’s jelly-derived MSCs (WJ-MSCs) exhibited increase in cell spread area and adhesion, with reduction in cellular migration under serum starvation stress. The changes in adhesion and migration characteristics were accompanied by formation of large number of super mature focal adhesions along with extensive stress fibres and altered ECM gene expression with notable induction in vitronectin (VTN) expression. NF-κβ was found to be a positive regulator of VTN expression while ERK pathway regulated it negatively. Molecular and phenotypic comparison studies with inhibition of these signalling pathways or knocking down of VTN under serum starvation established the correlation between increase in VTN expression and increased cellular adhesion with corresponding reduction in cell migration. VTN knockdown also resulted in reduction of super mature focal adhesions and extensive stress fibres, which were formed under serum starvation. Additionally, VTN induction was not detected in hypoxia-treated WJ-MSCs, and the MSCs showed no significant change in the adhesion or migration properties under hypoxia. However, when VTN expression was induced under hypoxia by ERK pathway inhibition, similar increase in cell spread area and adhesion were observed. Our study thus highlights VTN as a key player which is induced under serum starvation stress and possibly regulates the adhesion and migration properties of WJ-MSCs via focal adhesion signalling.

## Introduction

Mesenchymal stem cells (MSCs), being hypoimmunogenic, immunomodulatory and reparative in nature, have huge potential for clinical applications as a therapeutic candidate and personalised medicine for treatment of several inflammation-mediated and tissue injury related diseases [1]. The capability of MSCs to migrate and engraft at the site of injury followed by a balanced secretion of immunomodulatory and regenerative factors in response to the diseased microenvironment makes them a powerful candidate for cell-based therapy [2]. However, MSCs are unable to execute their full beneficial effect as the cells which are being cultured and expanded under favourable *in-vitro* culture conditions for clinical use, are not equipped to withstand the harsh *in-vivo* microenvironment prevalent at the recipient site due to inflammation or injury [3]. The diseased site is often marked by hostile microenvironmental conditions like ischemia, oxygen and nutrient deprivation, and presence of host-derived pro-inflammatory molecules, which could be detrimental to the viability and functionality of MSCs. The efficacy of MSC based cell therapy he nce gets compromised as transplanted MSCs, exhibit low migration rate and poor tissue engraftment at the site of disease and additionally, undergo significant loss in cell viability [4]. Migration, homing and adhesion are some of the key inherent properties of MSCs which play crucial role in strengthening their viability and functionality [5]. Thus, dissecting and understanding the basic regulation mechanisms underlying adhesion and migration of MSCs under stress conditions would help to overcome the current challenges and lead to better performing MSCs for clinical applications.

MSCs have been isolated from various adult as well as young tissue sources like bone marrow, adipose tissue, amniotic fluid, umbilical cord etc. via different methods of isolation, such as, enzymatic digestion, explant culture method etc. Depending on the tissue source or the method of isolation and culture, the therapeutic potential and functionality of MSCs may vary [6]. In fact, MSCs from different tissues, while exhibiting certain common basic characteristics, show some innate differences too. MSCs isolated from Wharton’s jelly (WJ), which is the perivascular region of umbilical cord, have been shown to possess better immuno-modulatory and regenerative properties along with higher proliferation capability as compared to many adult sources [7].

As MSCs are widely used for cell-based therapy towards ischemia-related tissue injury and other associated diseases which are marked by nutrient and oxygen deprived micro-environment, identifying the mode of regulation of cell adhesion and migration along with the underlying key factors under these stress conditions becomes pertinent. Extra-cellular matrix (ECM) proteins namely fibronectin, vitronectin (VTN) and collagen 1 act as motogenic factors both in soluble and insoluble forms and are known to play significant roles in recruitment of MSCs at the site injury for tissue repair and regeneration [8].

In our previous studies, we had shown that exposure of WJ-MSCs to serum starvation and ischemia-like condition *in-vitro* led to reduction in wound closure rate indicating impaired cell migration and in parallel, an ECM associated glycoprotein, VTN, also known as serum spreading factor, had shown up-regulation [9]. In fact, as some other groups reported in HEK239 cells, VTN regulated morphology and migration of cells via uPAR-matrix interaction [10]. VTN, also known for its adhesive role in tumor micro-environment, promoted tumor metastasis and regulated cell growth in neuroblastoma cells via cell-matrix interaction [11]. It was also demonstrated to interact with both integrin and uPAR via RGD and SMB domains, respectively, which increased the cell attachment by promoting focal adhesion (FA) component rearrangement at the FA sites [12].

Cell adhesion and cell migration usually display a biphasic relationship, whereby cell migration is maximum when cell-matrix adhesion is neither too high nor too low [13]. A recent study showed that inhibition of FA disassembly resulted in reduction of cell motility and flattening of MDA-MB-231 cells with elongated FAs formation and reduced rate of FA turnover [14]. Here, we observed that cell adhesion and migration of WJ-MSCs were altered on exposure to serum starvation condition *in-vitro* wherein cell adhesion increased with a corresponding decrease in cell migration. In parallel, an increase in VTN expression was noted in the serum-starved WJ-MSCs. By independently inhibiting ERK and NF-κβ signalling pathways, regulation of VTN expression was established by these two pathways. VTN silencing studies further confirmed its role in impacting altered morphology, adhesion and migration of serum-starved WJ-MSCs. VTN knockdown also led to reversal in expression of certain ECM genes like collagens along with reduction in expression of one of the focal adhesion (FA) complex proteins, vinculin. Again, assembly of large and elongated focal adhesions at the edge of prominent actin stress fibres in the central as well as peripheral regions of serum-starved WJ-MSCs, was abrogated with knockdown of VTN.

Physiological stress condition of ischemia is often marked by hypoxia as well, which is also prevalent in the natural in vivo niche of MSCs. Next, on subjecting WJ-MSCs to hypoxia stress condition *in-vitro*, interestingly, we noted that there was no detectable change in migration rate or adhesion of MSCs and the parameters were comparable to control WJ-MSCs, which corresponded well with the fact that there was no induction in the expression level of VTN, either. But, on inhibiting the ERK signalling pathway under hypoxia, a negative regulator of VTN expression, there was a marked increase in cell spread area and cellular adhesion along with induction in VTN expression.

Thus, our study establishes that VTN plays an important role in regulating the migration and adhesion of WJ-MSC under different stress conditions. This study, thus, could offer new strategies to positively modulate MSC migration and adhesion under physiological stress conditions, which could help improve their therapeutic efficacy following transplantation.

## Material and methods

### Cell culture

Human umbilical cords were collected after full term births (vaginal or cesarean delivery) with informed consent of the donor following the guidelines laid down by Institutional Ethics Committee (IEC) and Institutional Committee for Stem Cell Research and Therapy (IC-SCRT) at IISER, Kolkata, India. WJ-MSCs were isolated from the perivascular region of the umbilical cord termed as Wharton’s jelly by explant culture method as described previously [15]. All experiments were performed between passages 4 to 6 where cells were dissociated using TrypLE express (Life technologies), a gentle animal origin-free recombinant enzyme and seeded at a cell density of 5000 cells/cm^2^.

WJ-MSCs were initially cultured in complete medium comprising of KnockOut DMEM (Dulbecco’s modified Eagle’s medium) supplemented with 10% FBS, 2mM Glutamine and 1X PenStrep (all from Life technologies). For serum starvation stress, after 24h, cells were serum starved for the next 48h by replacing the medium with KnockOut DMEM medium supplemented with 2mM Glutamine and 1X PenStrep but no serum. For hypoxia treatment, cells were exposed to 2% oxygen for 48h and grown in complete medium comprising of KnockOut DMEM supplemented with 10% FBS, 2mM Glutamine and 1X PenStrep. Corresponding to the *in-vitro* serum starvation stress and hypoxia condition, the control cells were grown in complete medium for a total duration of 72h, in parallel. For different signalling pathway inhibition studies, cells were treated with specific small molecule inhibitors and their respective vehicle controls under serum starved condition or hypoxia for a period of 48h. ERK pathway inhibitor, FR-180204 and NF-κβ pathway inhibitor, BAY 11-7082 were added at an optimized concentration of 30µM and 4µM respectively (both from Sigma-Aldrich).

### siRNA transfection for knockdown studies

For siRNA mediated knockdown studies, WJ-MSCs were grown for 24-36h till they reached around 50-60% confluency. Transfection was performed using Lipofectamine 3000 in Opti-MEM I (both from Life Technologies) medium as per manufacturer’s protocol. For VTN and NF-κβ knockdown studies, endonuclease-treated siRNA pool (esiRNA) generated against VTN and p65 (all from Sigma-Aldrich) were used while for experimental control, mission universal negative control siRNA was used and all at a concentration of 30nM each. After 18-20h post transfection, WJ-MSCs were exposed to serum starvation for the next 36-48h and harvested as per the experimental requirement.

### Cell spread area calculation

For quantification of alteration in cell morphology, cell spread area of WJ-MSCs which were exposed to different experimental conditions was measured using ImageJ (NIH) software. Phase contrast imaging of WJ-MSCs was performed using Nikon Eclipse TS100 inverted microscope (Nikon) at 10X magnification. Cell spread area was calculated for 50 cells each from 3 independent biological samples and the data was plotted by fitting the data into Gaussian distribution using GraphPad Prism 8 software (GraphPad) to represent the cell spread area distribution across the population.

### De-adhesion dynamics

To study the alteration in adhesion property of WJ-MSCs under serum starved condition, cell de-adhesion dynamics was quantified by performing live cell time lapse imaging following treatment with 0.5mM EDTA (Sigma-Aldrich). For the experiment, live cells were treated with 0.5mM of EDTA and live cell imaging was performed using Olympus IX81 (Olympus) microscope to capture de-adhesion of the cells from the surface. Change in cell spread area across time was calculated and the normalized cell spread area was plotted against time. The data was fitted in Boltzmann’s equation in GraphPad Prism 8 software (GraphPad), from which two time constants were obtained which gave the quantitative measure of adhesion property of cells.

### Wound healing assay

To understand the effect of serum starvation on migration of WJ-MSCs, wound healing assay was performed. WJ-MSCs were seeded at a cell density of 6000 cells/cm^2^ and next, control or treated cells were scraped using a sterile micro-tip to create a uniform wound on the confluent monolayer of cells. Phase contrast imaging was performed at an interval of 2hs till wound closure using Olympus IX81 inverted microscope (Olympus) at 10X magnification and the micromanager software. The wound closure rate was quantified by measuring the change in the width of the wound across time using the ImageJ (NIH) software.

### Time lapse imaging

To further evaluate the migration pattern, directionality and migration rate of WJ-MSCs under serum starved condition, cells were plated at a cell density of 4000cells/cm^2^ in glass bottom dishes (SPL Lifesciences) and time lapse imaging was performed for a duration of 3h by capturing images at a frequency of 1 frame/min in confocal microscope LSM 710 (Axio Observer Z1, Carl Zeiss) equipped with an environmental chamber set at 37°C with 5% CO_2_. At least 50 cells per condition from three independent biological samples were analysed. The cell XY co-ordinates were obtained at each time point using ImageJ software (NIH) following which cell directionality, trajectory pattern and mean square displacement was quantified using DiPer software [16] and the cell migration rate was calculated using ZEN software (Zeiss).

### RNA isolation and cDNA synthesis

Total RNA was isolated using Trizol (Sigma-Aldrich) and the RNA yield was quantified using Nanodrop 2000 spectrophotometer (Thermo Scientific). cDNA synthesis was done using Verso cDNA synthesis kit (Thermo Scientific).

### Quantitative reverse transcription polymerase chain reaction

To quantify the mRNA expression of relevant genes, qRT-PCR reactions were done with PowerUp SYBR(tm) Green Master mix (Applied Biosystems) using ABI Biosystems 7500HT instrument (Applied Biosystems) or Biorad CFX96 Real Time System (Biorad). The fold change in mRNA expression was quantified by 2^-ΔΔCT^ method using compatible Applied Biosystems software. Validation of single, specific PCR products was done via melting curve analysis. *GAPDH* was considered as the endogenous control gene. The primer sequences and amplicon sizes are listed in Supplementary Table 1.

### Western Blotting

For quantification of protein expressed by WJ-MSCs under serum starvation, whole cell lysates were prepared using RIPA lysis buffer along with protease inhibitor cocktail (all from Santa Cruz Biotechnology). Protein concentration in whole cell lysate was quantified by Bradford assay. Polyacrylamide gel electrophoresis and western blotting were performed as per standard protocol. Primary and secondary antibodies used were anti-p65 (Santa Cruz Biotechnology), anti-VTN (Santa Cruz Biotechnology), anti-vinculin (Santa Cruz Biotechnology), anti-GAPDH (Santa Cruz Biotechnology) and HRP-linked anti-mouse IgG (Cell Signalling Technology).

### Immunofluorescence staining

To study the actin cytoskeleton arrangements and focal adhesion in WJ-MSCs on exposure to different treatments, phalloidin and paxillin immunofluorescence staining was performed as per standard protocol. The cells were grown on glass coverslips and exposed to specific treatment conditions as per the experiment. The primary and secondary antibodies used were anti-paxillin (Sigma-Aldrich or Genetex) and goat anti-mouse IgG H&L (Alexa Fluor® 568), goat anti-rabbit IgG (H+L) cross-adsorbed secondary antibodies (Alexa Fluor 568) (Thermo Scientific). Actin cytoskeleton was stained using Alexa Fluor(tm) 488 Phalloidin (Invitrogen), DAPI (Sigma-Aldrich) was used to stain the nucleus and then the coverslips were mounted on glass slides using VECTASHIELD anti-fade mounting medium. The phalloidin images were captured using ZEN software (Zeiss) at 40X magnification in Zeiss Apotome module microscope and the paxillin phalloidin co-immunostaining images were captured in Leica SP8 confocal microscope using oil immersion 63X objective and deconvoluted using Leica Lightening software.

### Statistical Analysis

All data are presented as mean ± standard error of the mean (SEM) from at least 3 independent biological samples. Data analysis and graphical representations were performed using GraphPad Prism 5/ 8 software (GraphPad). Statistical comparisons were assessed using the two-tailed Student t-test, one-way ANOVA or two-way ANOVA. Significance was accepted at *p* ≤ 0.05.

## Results

### Serum starvation impaired the migration ability of WJ-MSCs

As reported previously by our group [17], exposure to serum starved condition resulted in flattened morphology with significant increase in cell spread area (data not shown) and cellular adhesion in WJ-MSCs (Fig. 1A, B). For the de-adhesion assay, a minor modification was introduced here where WJ-MSCs were treated with warm EDTA only. A slower de-adhesion was reflected by the right shift of the sigmoidal curve (Fig. 1A) along with increase in τ1 and τ2 values from 15.48±0.93s to 79.5±5.85s and 7.38±0.56s to 30.52±1.67s (Fig. 1B, *p* < 0.001), respectively. In addition to these phenotypic changes, next we sought to investigate the effect of serum starvation stress on cell migration of WJ-MSCs.

**Figure 1:**
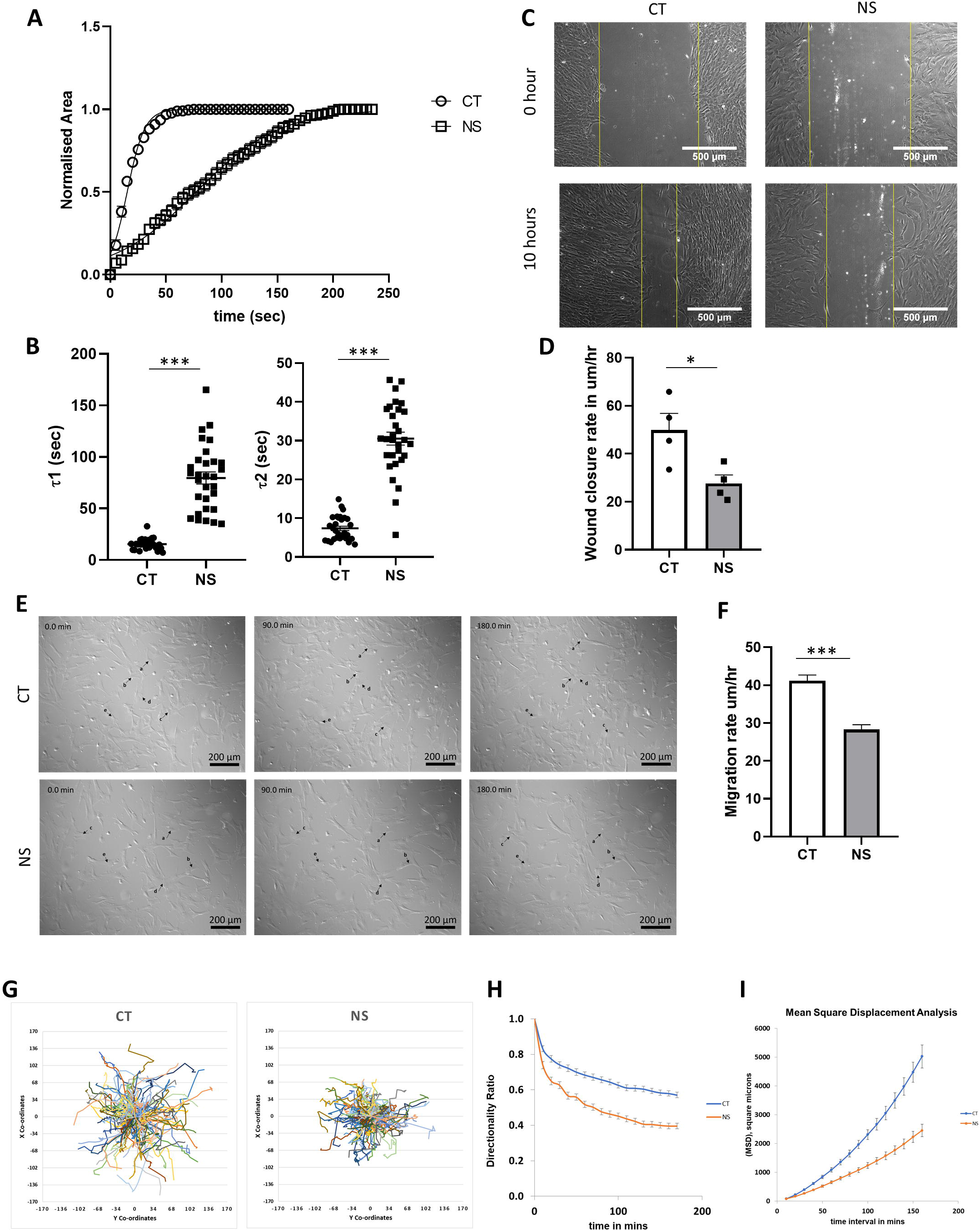
De-adhesion dynamics of individual WJ-MSCs on treatment with EDTA was compared between control and serum-deprivation condition. Effect of serum deprivation stress on time constants of retraction was recorded with the help of time lapse imaging. The normalized area-vs.-time data were fitted to a Boltzmann sigmoid equation to determine the time constants τ1 and τ2 (A). Delayed de-adhesion under serum deprivation led to significant differences in τ1 and τ2 between serum-deprived and control WJ-MSCs (B). Comparison of wound induced migration between control and serum-deprived WJ-MSCs. Confluent monolayer cultures of control and serum-deprived WJ-MSCs were scratched with a sterile pipette tip at 0h. Representative images of migration post scratch assay were obtained at 10h, Scale bar: 500µm (C). Wound closure rate was also quantified. Each bar represents mean ± SEM (n = 5) (D). Representative DIC still images were captured from the time-lapse videos of control and serum-deprived WJ-MSCs. Black arrows identify the migration of some randomly selected cells. Numbers on the images indicate time in minutes and results are representative of three independent biological samples (n = 3) (E). To further investigate in detail the impact of serum deprivation on migration of WJ-MSCs, migration speed (F), trajectories of cells (G), directionality ratio (H) and MSD of cell body (H) were calculated and compared between control and serum-deprived WJ-MSCs for n = 150 cells from three different biological samples. Serum-deprived condition is denoted as no serum (NS) in the figure. Each bar represents mean ± SEM. (\**p*< 0.05, \*\*\**p*< 0.001).

First, cell migration was assayed via *in-vitro* scratch-induced wound healing assay (Fig. 1C). Cell migration rate, assessed during wound closure, indicated that serum-starved WJ-MSCs exhibited slower wound closure rate (27.6±3.5µm/h) as compared to control WJ-MSCs (49.9±6.8µm/h) (Fig. 1D; *p <* 0.05).

To analyse and gain additional insights, we investigated the impact of serum starvation on the migration pattern of WJ-MSCs, and live cell time lapse imaging was performed (Fig. 1E, Supp. Video 1-2). Cell migration was quantified in terms of single cell migration rate (Fig. 1F, *p* < 0.001), single cell trajectory pattern (Fig. 1G), directionality ratio (Fig. 1H) and mean square displacement (Fig. 1I) using ZEN and DiPer software [16]. Both single cell migration rate and mean square displacement data indicated that the cell migration decreased significantly on exposure to serum starvation stress as compared to the control condition (Fig. 1F, 1I). Corresponding to the change in migration rate, the single cell trajectory pattern showed lower spread area indicating impaired migration of serum-starved WJ-MSCs (Fig. 1G), along with loss of directionality as also suggested by the directionality ratio plot (Fig. 1H).

### Expression and regulation of VTN in WJ-MSCs under serum starvation stress

Next, we investigated the expression and regulation of VTN under serum starved condition in co-relation with the phenotypic changes observed. In our previous studies [17,18], we had observed that expression of VTN, an ECM component known to be responsible for cell adhesion, was up-regulated under both serum starvation and febrile temperature stress conditions and this was accompanied by increase in cell spread area and cell adhesion. In the present study under serum starved condition, VTN expression again exhibited a strong upregulation both at mRNA and protein levels, though not significant (Fig. 2A, B). Next, to determine the signalling pathways involved in regulating the expression of *VTN* at the mRNA level, WJ-MSCs were treated with different small molecule inhibitors for different signalling pathways (Fig. 2C, Supp. Fig. 1A). The data identified ERK and NF-κβ signalling pathways being involved in regulating VTN expression.

**Figure 2.**
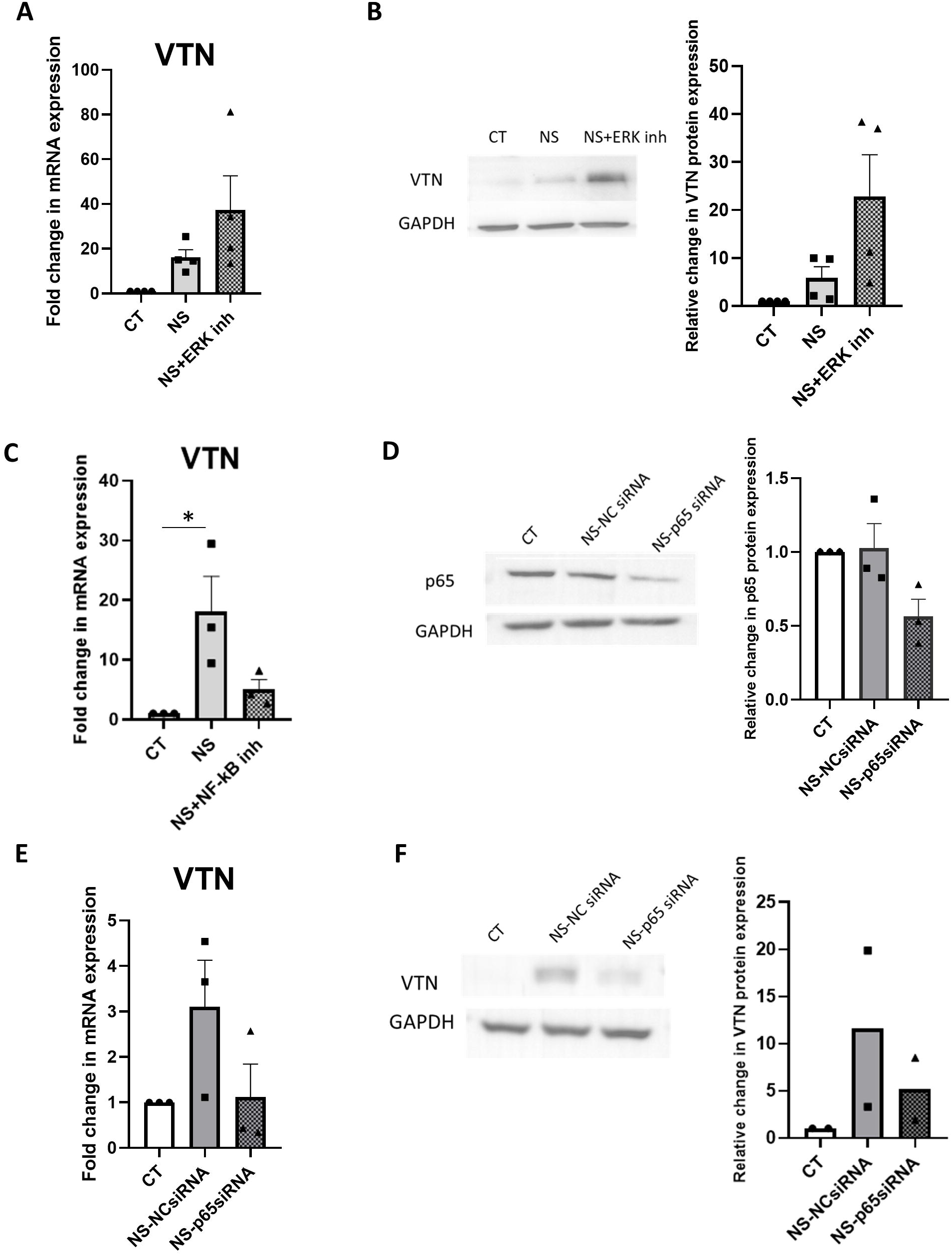
Effect of ERK pathway inhibition on VTN expression in serum-deprived WJ-MSCs. The mRNA and protein expression levels of VTN were detected by qRT-PCR (n=3) (A) and Western blotting (n=3) (B), respectively, in WJ-MSCs treated without and with ERK pathway inhibitor for 48h under serum deprivation condition. Impact of NF-κβ pathway inhibition on expression of VTN. WJ-MSCs were treated with BAY 11-7082, an inhibitor of NF-κβ pathway or transfected with a p65-targeted esiRNA for 48h under serum-deprived state. The reduction in p65 protein level on transfecting with p65-targeted esiRNA was confirmed by Western blotting (D). The mRNA and protein expression levels of VTN were detected by qRT-PCR (n=3) (C, E) and Western blotting (n=2) (F), respectively. *GAPDH* was used as endogenous control in qRT-PCR as well as a loading control in Western blotting. Band densities of Western blotting were quantified relative to GAPDH. Data shown are representative of at least two independent biological samples. Serum-deprived condition is denoted as no serum (NS) in the figure. Each bar represents mean±SEM (\**p*< 0.05, \*\**p*< 0.01, \*\*\**p*< 0.001).

Inhibition of ERK pathway by small molecule inhibitor, FR180204 under serum starved condition further increased the expression of VTN both at mRNA and protein levels as compared to only serum starved condition, though not significant, indicating that ERK signalling pathway negatively regulated the expression of VTN (Fig. 2A, B), while NF-κβ pathway inhibition by small molecule inhibitor, BAY 11-7082, reduced *VTN* mRNA expression (Fig. 2C) (not significant).Further, siRNA mediated knockdown of NF-κβ subunit, p65, (Fig. 2D) demonstrated that NF-κβ signalling pathway was a positive regulator of VTN expression, since its inhibition reversed the induced expression of VTN under serum starved condition at both mRNA and protein levels (Fig. 2E, F) (not significant).

### Impact of ERK and NF-κβ pathway silencing on adhesion and migration behaviour of WJ-MSCs under serum starvation stress

To validate whether VTN could be responsible for causing the increase in cell spread area and cell adhesion, and decrease in cell migration of WJ-MSCs under serum starvation, we analysed these parameters under ERK or NF-κβ pathway inhibition.

It was observed that cell spread area, along with cell adhesion exhibited a further increase on inhibiting ERK pathway under serum starved condition (Fig. 3A-D; *p <* 0.001). The cell adherence was quantified in terms of de-adhesion dynamics of the cell where cell adhesions were severed by EDTA treatment and cellular detachment phenomena was analysed as decrease in cell spread area with respect to time. The further increase in τ1 and τ2 values from 59.32±3.7s to 93.69±5.5s and 24.63±1.28s to 38.87±1.96s, respectively, clearly indicated that WJ-MSCs adhered more strongly on inhibiting ERK signalling pathway under serum starvation as compared to without ERK pathway inhibition (Fig. 3D). Also, ERK pathway inhibition under serum starved condition led to stronger reduction in cellular migration as compared to only serum starved condition, where the wound closure rate reduced from 31.54±1.89µm/h to 15.12±2.64µm/h as observed by *in-vitro* wound healing assay (Fig. 3E, F; *p* < 0.05).

**Figure 3:**
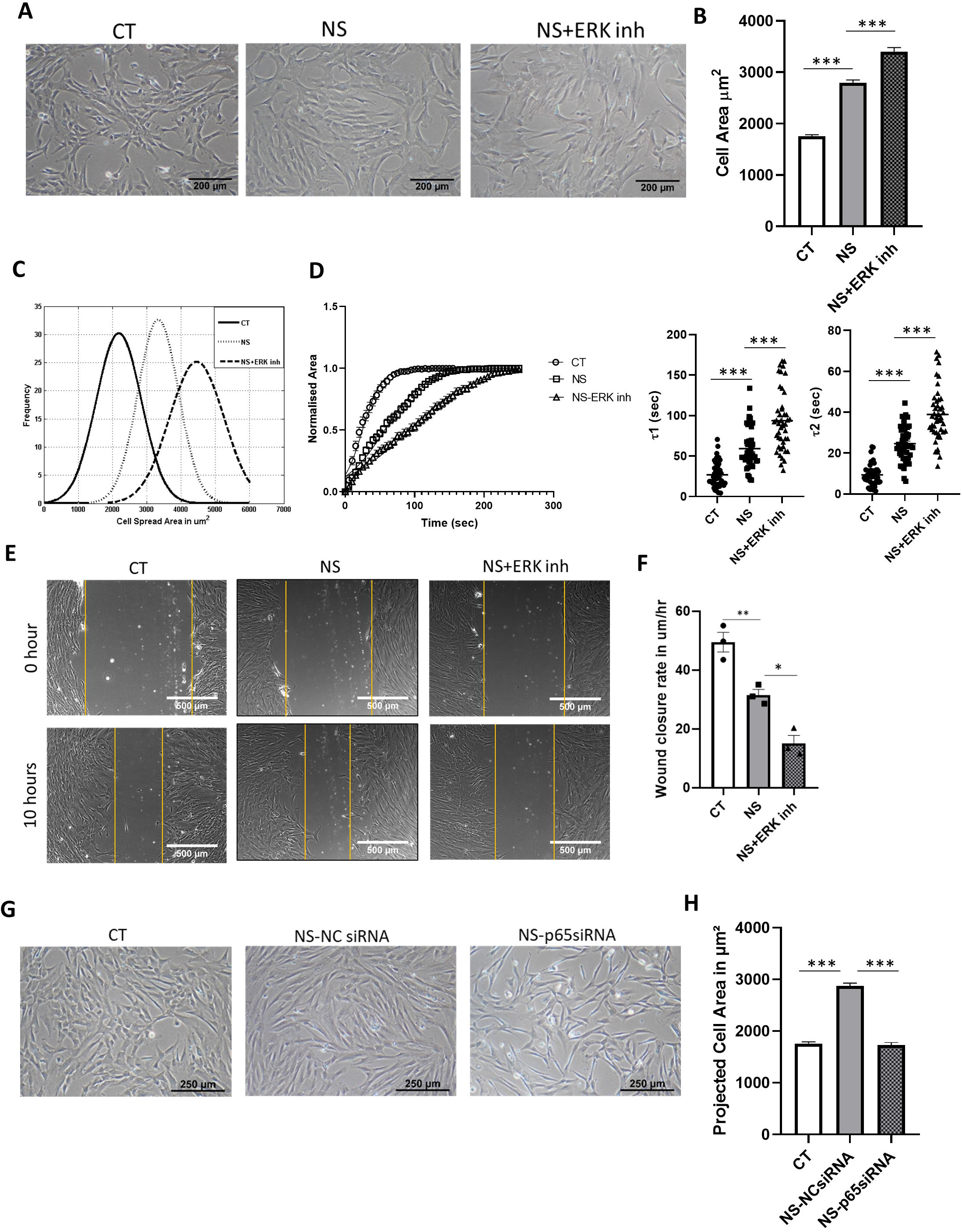
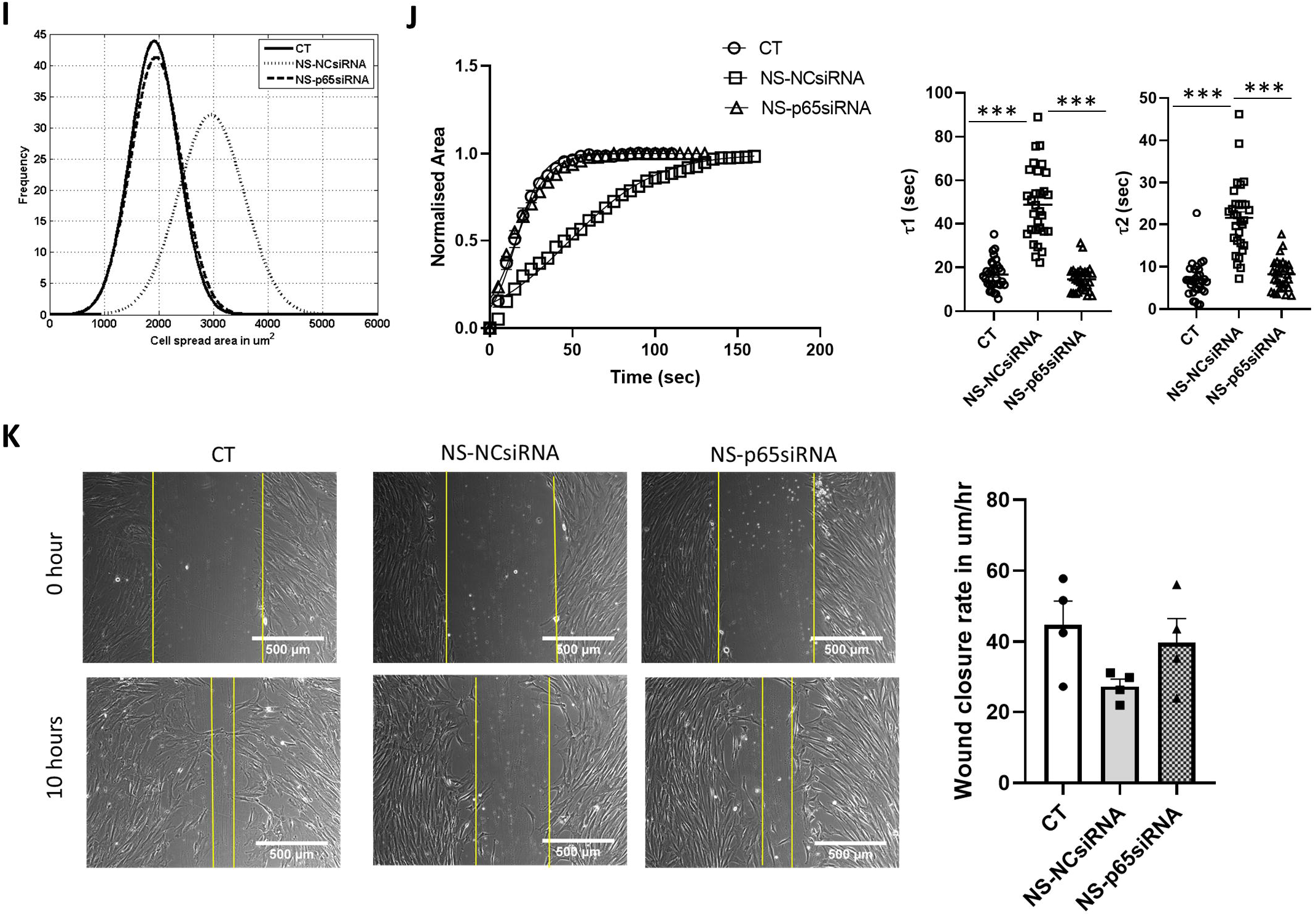
Assessing the impact of ERK pathway inhibition on cell spread area, de-adhesion dynamics and migration of serum-deprived WJ-MSCs. WJ-MSCs were treated with FR180204 for 48h under serum deprived condition. Representative phase contrast images showing morphology are displayed (n = 3) (A). Cell area was quantified and compared for 150 cells from 3 different biological samples (B). A representative Gaussian distribution plot from 50 cells is presented to compare cell area between control and serum-deprived WJ-MSCs without and with ERK inhibitor treatment (C). De-adhesion dynamics of FR180204 treated WJ-MSCs under serum deprivation. With the help of time lapse imaging, cell shape changes during de-adhesion were quantified and recorded. The normalized area-vs.-time data were fitted to a Boltzmann sigmoid equation to determine the time constants τ1 and τ2 (n=3) (D). Comparison of scratch-induced migration between control and serum-deprived WJ-MSCs without and with FR180204. Confluent monolayer cultures of control and treated WJ-MSC were wounded with a sterile pipette tip at 0h. Images of migration post scratch assay were obtained at 10h. Scale bar: 500μm. Representative images from three independent biological samples (n=3) are exhibited (E). Wound closure rate was calculated and compared between control and treated WJ-MSCs. Bar represents mean ± SEM (n=3) (F). Effect of NF-κβ pathway inhibition on cell spread area, de-adhesion dynamics and migration of serum-deprived WJ-MSCs. WJ-MSCs were transfected with NC siRNA or p65-targeted esiRNA under serum-deprived condition. Representative phase contrast images showing morphology are displayed (n=3) (G). Cell area was calculated for WJ-MSCs transfected with p65-targeted esiRNA or NC siRNA under serum starvation and plotted against control WJ-MSC samples. A sample size of 150 total cells, from three different biological samples, was used (H). A representative Gaussian distribution plot for 50 cells is presented to compare the cell area between control and transfected WJ-MSCs under serum deprivation (I). De-adhesion dynamics of p65 esiRNA transfected WJ-MSCs under serum deprivation. The change in cell area across time was quantified and fitted in Boltzmann sigmoid equation to determine the time constants. Differences in τ1 and τ2 between negative control siRNA and p65 esiRNA treated cells were plotted (J). Comparison of scratch wound induced migration between control WJ-MSCs and WJ-MSCs transfected with NC siRNA or p65-targeted esiRNA under serum starvation was assessed. Representative images are shown (n = 4). Wound closure rate in μm/h was determined from scratch induced wound healing assay (K). Serum-deprived condition is denoted as no serum (NS) in the figure. Each bar represents mean ± SEM. (\**p*< 0.05, \*\**p*< 0.01, \*\*\**p*< 0.001).

In contrast, NF-κβ pathway silencing via siRNA mediated knockdown of p65 led to reduction in cell spread area and cellular adhesion (Fig. 3G-J). Faster de-adhesion was assessed from left shift of the sigmoidal curve and also, the τ1 and τ2 graphs showed that knockdown of p65 led to reduction in time constant values from 49.15±2.5s to 16.43±0.97s and 23.2±1.28s to 8.68±0.5s, respectively, as compared to negative control siRNA treated WJ-MSCs under serum starved condition (Fig. 3J; *p* < 0.001). Further, the reduction in cell migration also got rescued with p65 knockdown under serum starved condition, though not significant, as evident from increase in the wound closure rate shown in the *in-vitro* wound healing assay data (Fig. 3K).

Overall, we observed that increase in VTN expression corresponded to increase in cell spread area and cellular adhesion along with reduction in cellular migration.

### VTN regulated the morphology, adhesion and migration of WJ-MSCs under serum starved condition

To specifically confirm the role of VTN in regulation of adhesion and migration of WJ-MSCs under serum starved condition, we analysed the effect of siRNA mediated silencing of VTN on these parameters. We observed that siRNA mediated knockdown of VTN (Fig. 4A) (not significant) under serum starvation led to distinct reversal in cell morphology (Fig. 4B, C; *p* < 0.001) and cell adhesion (Fig. 4D; *p* < 0.001) as compared to NC siRNA treated WJ-MSCs. Knockdown of VTN resulted in decrease in τ1 and τ2 values from 48.7±2.29s to 14.23±0.97s and 23.05±1.2 to 7.4±0.5s, respectively, as obtained from the sigmoidal curve of de-adhesion dynamics indicating that VTN did play an important role in promoting cellular adhesion under serum starved condition (Fig. 4D). Moreover, impaired migration of WJ-MSCs under serum starved condition also got rescued post VTN knockdown as evident from improved single cell trajectory pattern and increased single cell migration rate (Fig. 4E-G, Supp. Video 3-5, *p* < 0.001). Along with the recovery in migration pattern, the directionality ratio and the mean square displacement of cells also got rescued with VTN knockdown under serum starved condition (Fig. 4H, I).

**Figure 4:**
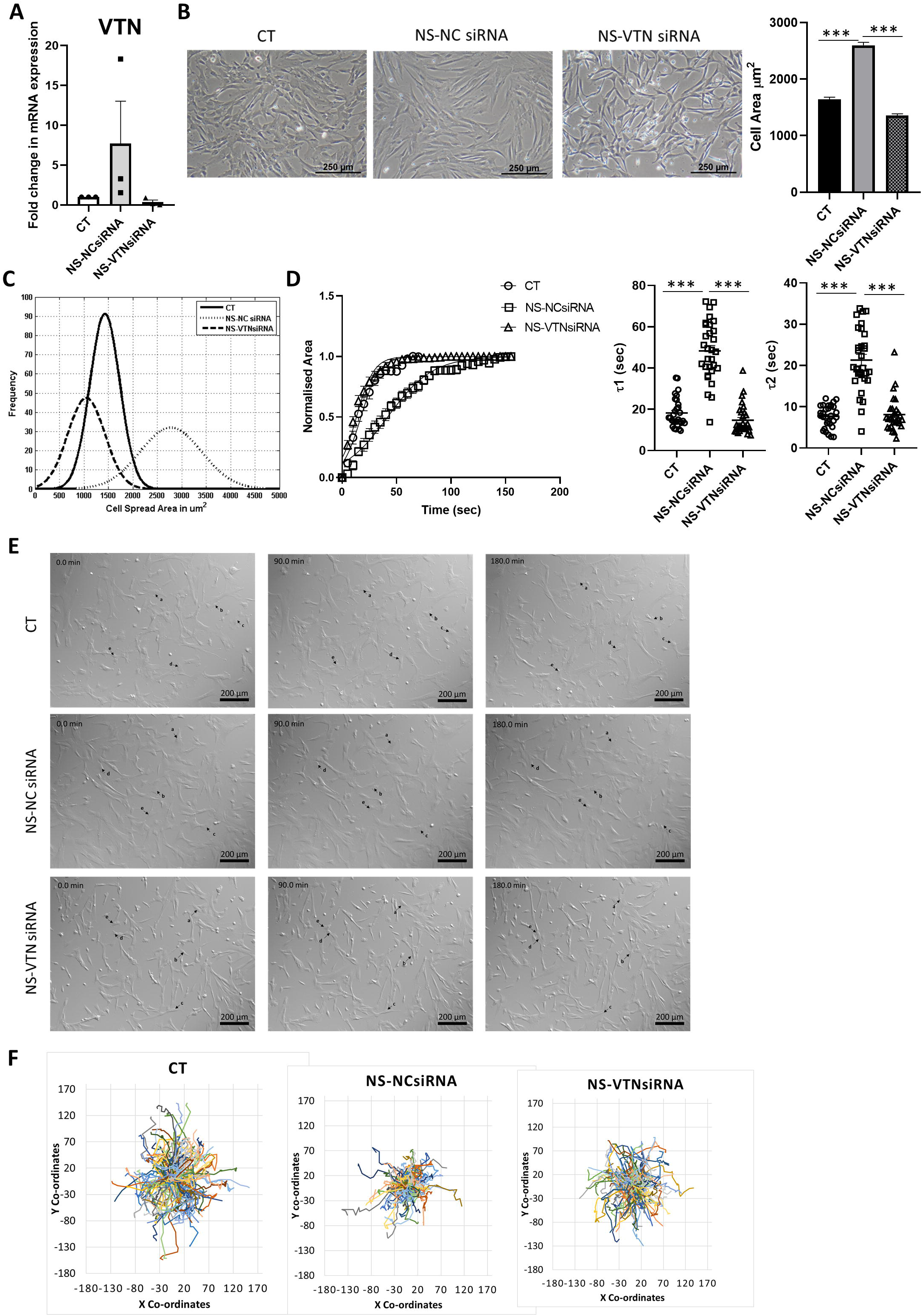
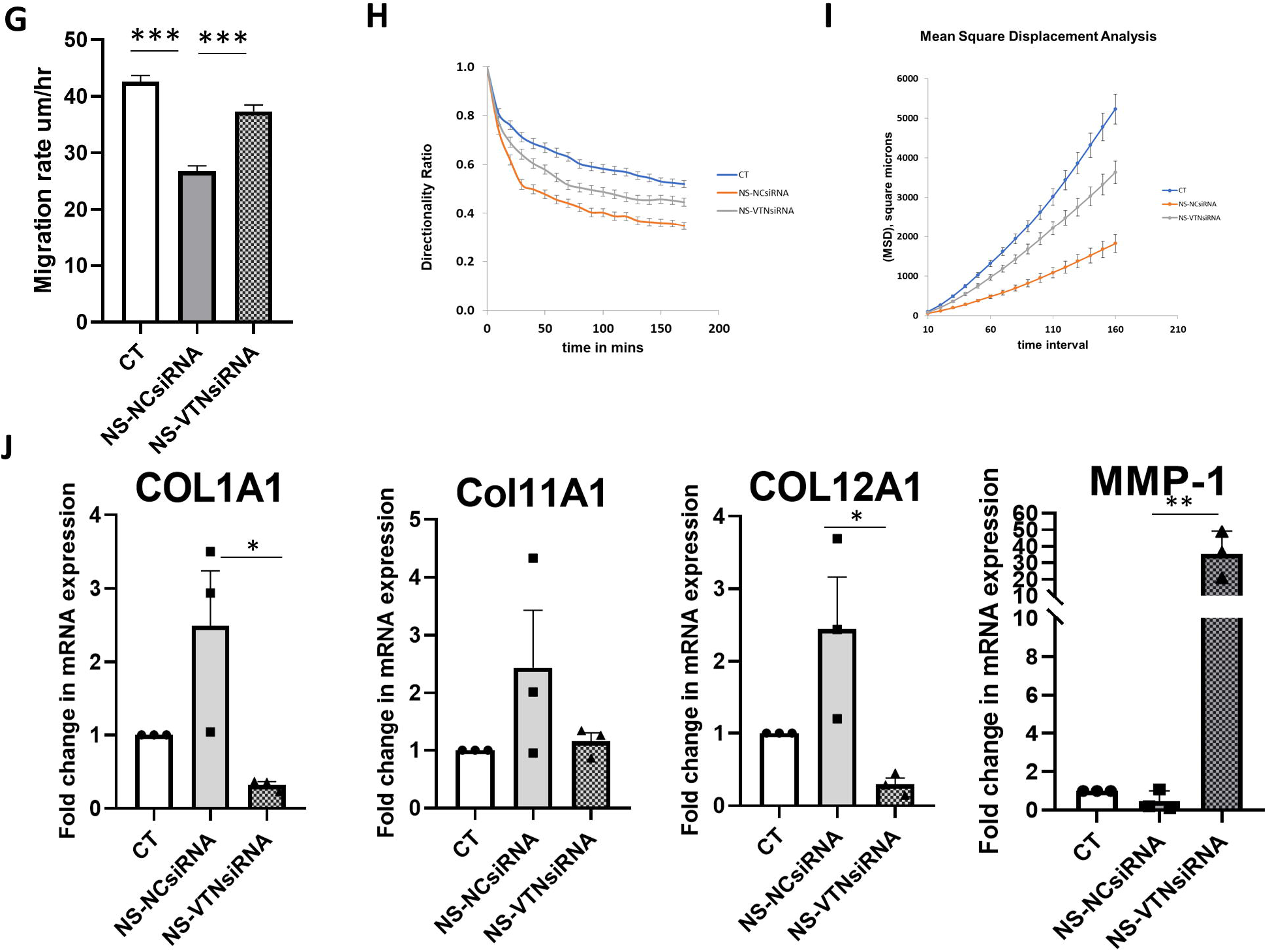
Effect of VTN knockdown on morphology, adhesion and migration of WJ-MSCs. Knockdown of VTN in WJ-MSCs, following transfection with VTN esiRNA, was validated by qRT-PCR. GAPDH was used as house-keeping gene (n=3) (A). Cell morphology images of WJ-MSCs transfected with VTN esiRNA or NC siRNA under serum starved condition for 48h in comparison to control condition are displayed. Scale bar: 500µm. Cell spread area was compared between control WJ-MSCs and WJ-MSCs transfected with VTN esiRNA or NC siRNA under serum starved condition. A total of 150 cells from three different biological samples were used (B). A representative Gaussian distribution plot from 50 cells is shown (C). Effect of VTN knockdown on de-adhesion dynamics of serum-starved WJ-MSCs was quantified in terms of time taken for cellular detachment following EDTA treatment. The normalized change in cell spread area vs time was fitted to Boltzmann sigmoidal equation to obtain the time constants, τ1 and τ2. Significant reversal in the time constant values, τ1 and τ2 with knockdown of VTN under serum starved condition was observed. A total of 45 cells from 3 different biological samples were used for the assay (D). Impact of VTN knockdown on migration of serum-starved WJ-MSCs. Representative DIC images from different time intervals are marked with small black arrows to highlight migration of randomly selected cells from time-lapse imaging (E). Comparison of migration rate of control vs WJ-MSCs transfected with VTN esiRNA or NC siRNA under serum starved condition is shown by the bar graph for 150 cells from three independent biological samples (G). For further insight, cell trajectory pattern (F), directionality ratio(H) and mean square displacement (I) were quantified from the analysis of DIC time lapse imaging videos for 150 cells from three independent biological sample. mRNA expression levels of ECM genes like *MMP-1, COL1A1, COL11A1 and COL12A1* was quantified following VTN knockdown under serum starved condition using qRT-PCR (n=3) (J). Serum-deprived condition is denoted as no serum (NS) in the figure. Each bar represents mean ± SEM. (\**p*< 0.05, \*\**p*< 0.01, \*\*\**p*< 0.001).

ECM components like collagens and MMPs are known to play important functional roles in cell adhesion, migration, differentiation, regeneration etc [19]. On investigating further, increased expression was noted for several collagen molecules, *COL IA, COL 12A, COL 11A* under serum starvation condition, which was reverted when VTN was knocked down, as presented in Fig. 4J. Meanwhile, *MMP-1*, an ECM enzyme responsible for protein degradation and known to aid in cell migration, was downregulated under serum starved condition. However, on knocking down VTN, *MMP-1* expression got strongly up-regulated at mRNA level (Fig. 4J; *p* < 0.05). This indicated that VTN could be involved in regulating the expression of ECM genes like collagens and MMPs, as a consequence of which the adhesion and migration parameters of WJ-MSCs got altered under serum starved condition.

### VTN mediated increase in focal adhesions responsible for the impaired migration under serum starvation stress

Cell migration comprises of a series of transitions which are co-ordinately regulated by cellular adhesions and actin cytoskeleton of a cell [20]. Previous literature documented that massive stress fibre formation reduced their dynamics resulting in defects in cell motility [21]. To understand the mechanism underlying the dysregulation in cellular adhesion and migration in WJ-MSCs under serum starvation stress, phalloidin and paxillin immunofluorescence staining was performed. Phalloidin immunofluorescence staining indicated that serum starvation also resulted in robust increase in F-actin in the form of stress fibres in WJ-MSCs, which underwent reversal with VTN knockdown. Quantification of mean intensity of phalloidin staining confirmed the same (Fig 5A; *p <* 0.001).

**Figure 5:**
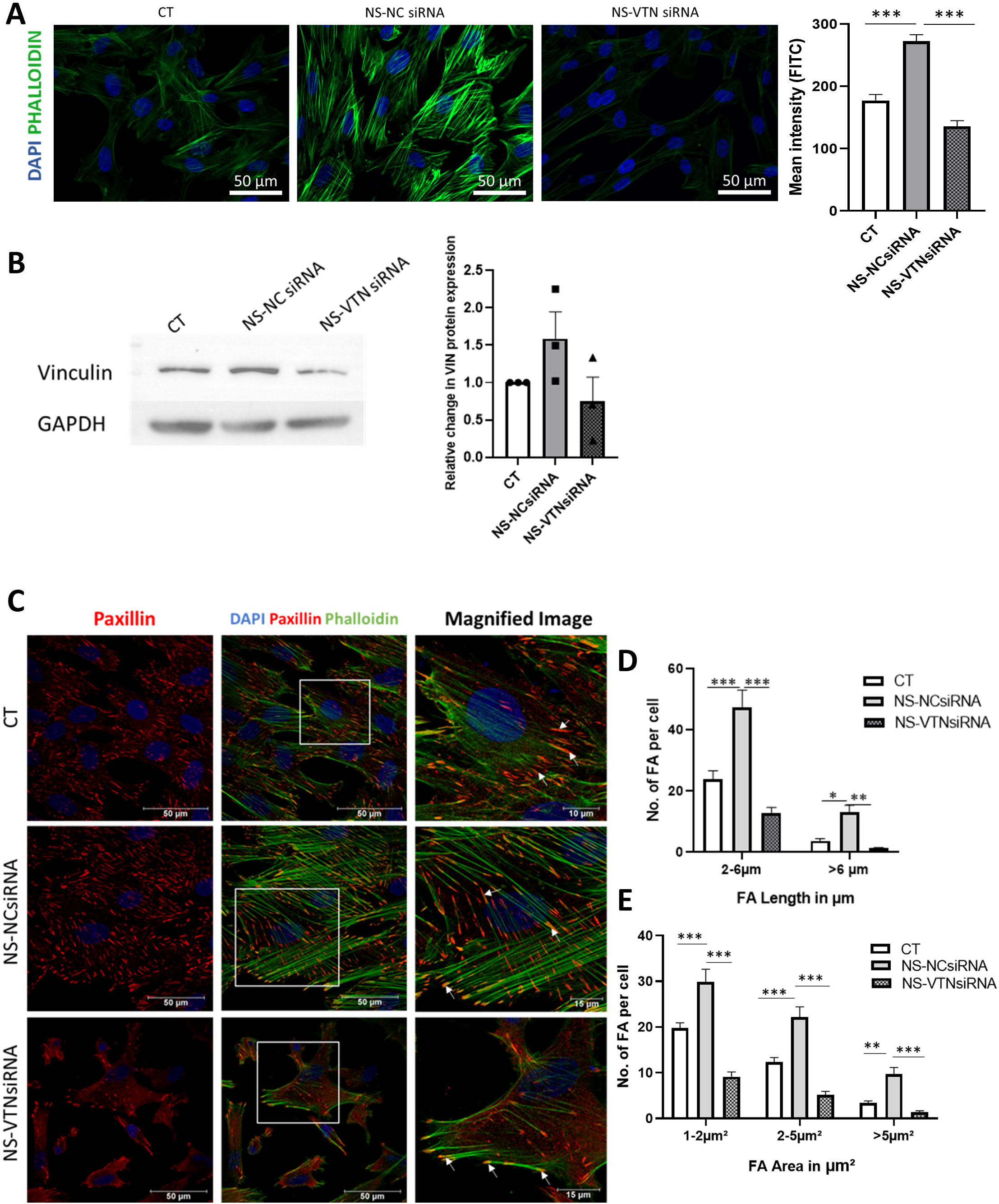
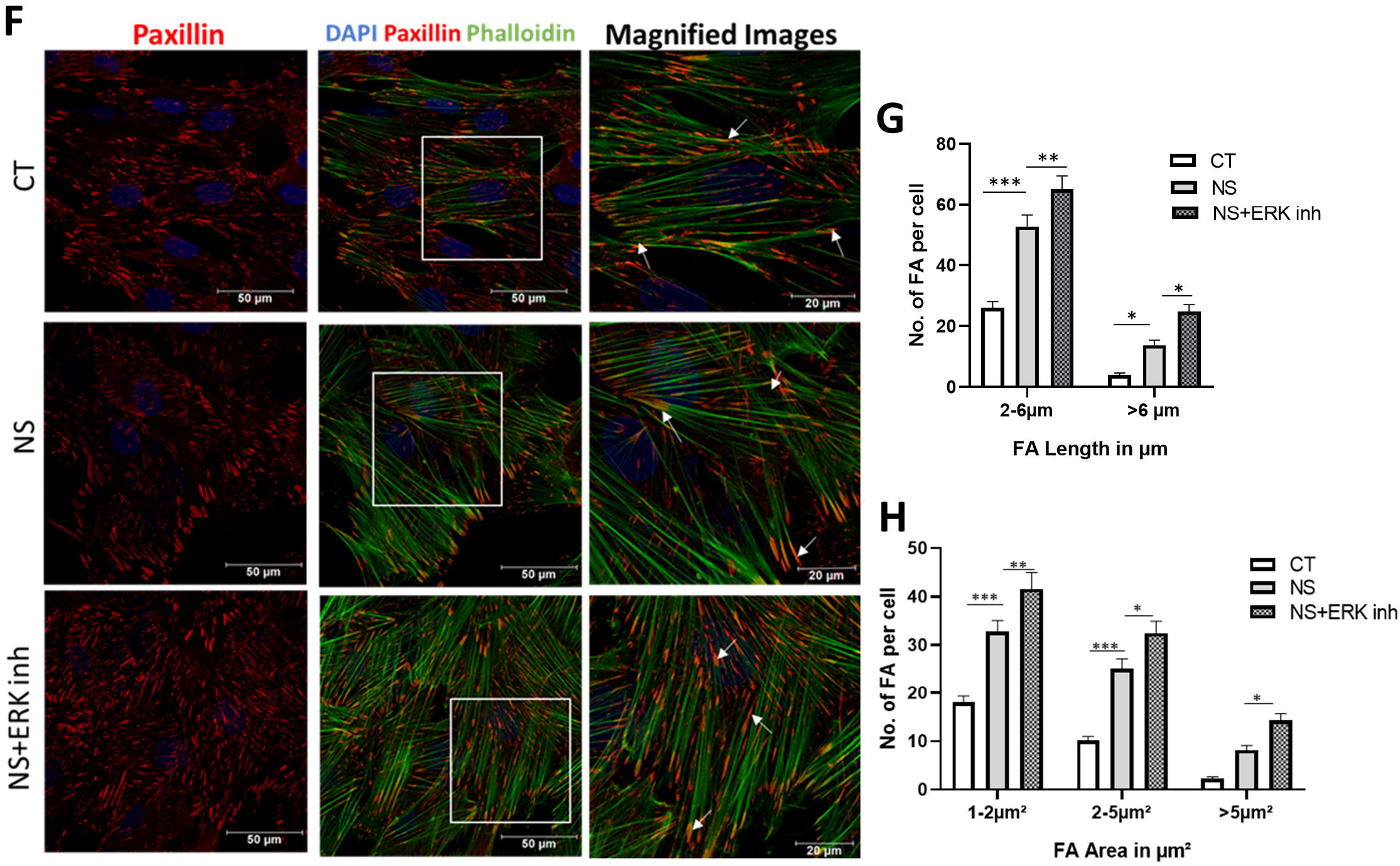
VTN dependent alteration in actin cytoskeleton and focal adhesion pattern. To compare the changes in actin cytoskeleton organisation between control WJ-MSCs and WJ-MSCs transfected with VTN esiRNA or NC siRNA, actin stress fibres were visualized via immunofluorescence using Alexa Fluor 488-phalloidin (green) while DAPI (blue) was used to stain nuclei. Reduction in actin stress fibre formation upon VTN knockdown under serum starved condition is demonstrated in the representative immunofluorescence images (40X magnification) (n=3). Scale: 50µm. Mean intensity of phalloidin staining was quantified and plotted (A). The protein expression of vinculin under VTN knockdown condition was detected by Western blotting experiment. Representative blot images are displayed. Band densities of Western blots were quantified relative to GAPDH. Data shown are representative of at least three independent biological samples (n=3) (B). Representative paxillin and phalloidin co-immunostaining images of WJ-MSCs are displayed for comparison of focal adhesion pattern under serum starvation with or without VTN knockdown. Paxillin staining of focal adhesions was performed using Alexa Flour 568 (red) labelled antibody and actin filaments were stained with Alexa Fluor 488-phalloidin (green). Scale: 50µm. The increased number of FAs under serum starvation stress was reduced following VTN knockdown. Along with increased FA numbers under serum starvation, distribution of FAs were observed all across the cell including the cell periphery, while under control or VTN knockdown conditions the cell periphery got predominantly stained for FAs (white arrows) (C). FA length was quantified in terms of classical FA (2-6μm) and super mature FA (>6μm) and FA area was quantified and grouped (1-2μm^2^, 2-5μm^2^ or >5μm^2^) from at least 100 cells for each sample. Number of FAs per cell was plotted from three independent biological samples (n=3) (D, E). Representative paxillin and phalloidin co-immunostaining images are displayed for comparison of focal adhesion pattern under serum starvation with or without ERK pathway inhibition. Paxillin staining of focal adhesions was done using Alexa Flour 568 (red) labelled antibody and actin filaments were stained with Alexa Fluor 488-phalloidin (green). Scale: 50µm. ERK pathway inhibition further increased the number of FAs, with prominent distribution of larger FAs across the cell compared to serum starvation condition (F). FA length and FA area were quantified from at least 100 cells from two independent biological samples (n=2) and the number of FAs per cell was plotted (G, H). Each bar represents mean ± SEM (\**p*< 0.05, \*\**p*< 0.01, \*\*\**p*< 0.001).

During migration, the stress fibres together with focal adhesions (FAs) at their ends constitute a major mechano-sensing machinery in the cells [22]. FAs are multi-protein structures connecting the intracellular cytoskeleton to the ECM. In our study, protein expression of vinculin, one of the FA components, was found to be elevated under serum starvation, but with VTN knockdown this increase in expression was abrogated (Fig. 5B, not significant). Previous reports showed that fast moving cells had dot-like nascent FAs while slow migrating cells possessed large streak like FAs [23]. The length of classical FAs was defined at 2-6μm, while the super mature FA length ranged between 6-30μm [24]. Thus, morphometric properties of FA like length and area were quantified in serum-starved WJ-MSCs, under different treatment conditions. The paxillin immunostaining (Fig. 5C) highlighted that the FAs formed in control WJ-MSCs are small and majorly located at the periphery of the cell. While, under serum starvation, FAs were majorly large streak like in appearance and located not only at the periphery but also across the cell, marking the distal ends of the extensive stress fibres (marked with white arrows) attaching the cytoskeleton with the ECM substrate (Fig. 5C). Knockdown of VTN under serum starvation abrogated this change in focal adhesion and cytoskeleton arrangement. Quantification of FA lengths showed presence of significantly higher number of super mature FAs of >6µm (from 3.77±0.63 to 13.1± 2.28) and classical FAs of 2-6μm (from 23.96± 2.67 to 47.25±5.78) per cell under serum starvation as compared to control condition. Knocking down of VTN under serum starvation significantly reduced the number of both classical FAs (from 47.25 ± 5.78 to 12.75±1.93) and super mature FAs (from 13.1±2.28 to 1.3±0.23 (Fig. 5D, *p <* 0.001), as compared to only serum starvation stress. Further, FA area quantification data corroborated the same (Fig. 5E, *p <* 0.001).

As we had noted stronger upregulation in VTN expression along with reduction in cell migration with ERK pathway inhibition under serum starvation condition, to further validate, paxillin immunostaining was performed under serum starvation condition in the presence of ERK pathway inhibitor. Immunostaining data showed the presence of more extensive stress fibres marked by array of elongated FAs all across the cells (marked with white arrows) under ERK pathway inhibition as compared to only serum starved condition (Fig. 5F). Morphometric analysis of length of focal adhesions showed further increase in number of classical FAs (from 52.76± 3.72 to 65.23± 4.31) and super mature FAs (from.13.74±1.7 to 24.79±2.33) compared to only serum starvation treatment (Fig. 5G, *p <* 0.001). FA area quantification data corroborated the same (Fig. 5H, *p <* 0.001). Collectively, the possible involvement of VTN in inducing focal adhesion formation, which in turn, impacted the cellular adhesion and migration of WJ-MSCs under certain defined stress conditions, was highlighted.

### Changes in morphology, adhesion and migration of WJ-MSCs under hypoxia condition in response to VTN expression level

A hypoxic microenvironment is associated with physiological stem cell niche and is known to support and improve survival and migration of MSCs [25]. Thus, to compare against serum starvation stress where the adhesion and migration of WJ-MSCs got adversely affected, we next exposed WJ-MSCs to a hypoxic milieu of 2% O_2_ and studied the expression of VTN, and examined the adhesion and migration of WJ-MSCs to further validate the role of VTN. Interestingly, under hypoxia condition, WJ-MSCs assumed small spindle shaped morphology (Fig. 6A-C, *p <* 0.01) with reduced adhesion as compared to control condition (Fig. 6D, *p <* 0.001). The cell trajectory pattern and the directionality ratio also mostly remained unaltered under hypoxia with a small increase in migration rate (Fig. 6E-G). Corresponding to these findings on the adhesion/migration characteristics, VTN expression level did not show any significant change under hypoxia treatment. (Fig. 6H, I). Phalloidin staining of hypoxia exposed WJ-MSCs did not indicate any increase in accumulation of actin stress fibres either, and the overall mean intensity of phalloidin staining remained similar between control and hypoxic-WJ-MSCs (Fig. 6J). Thus, the alterations witnessed in adhesion/migration parameters of WJ-MSCs under serum starvation were absent in hypoxic WJ-MSCs and there was neither any induction in VTN expression.

**Figure 6:**
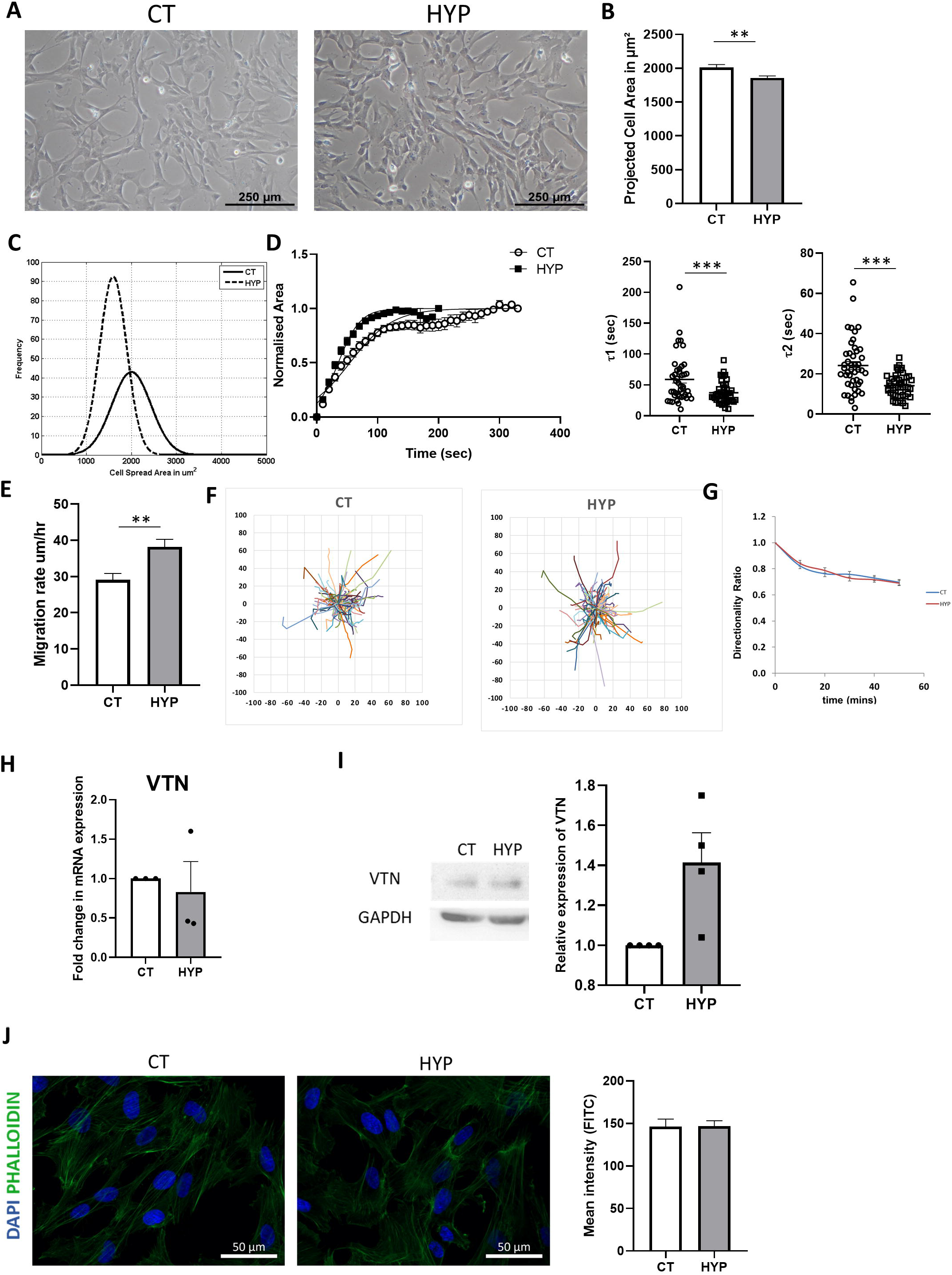
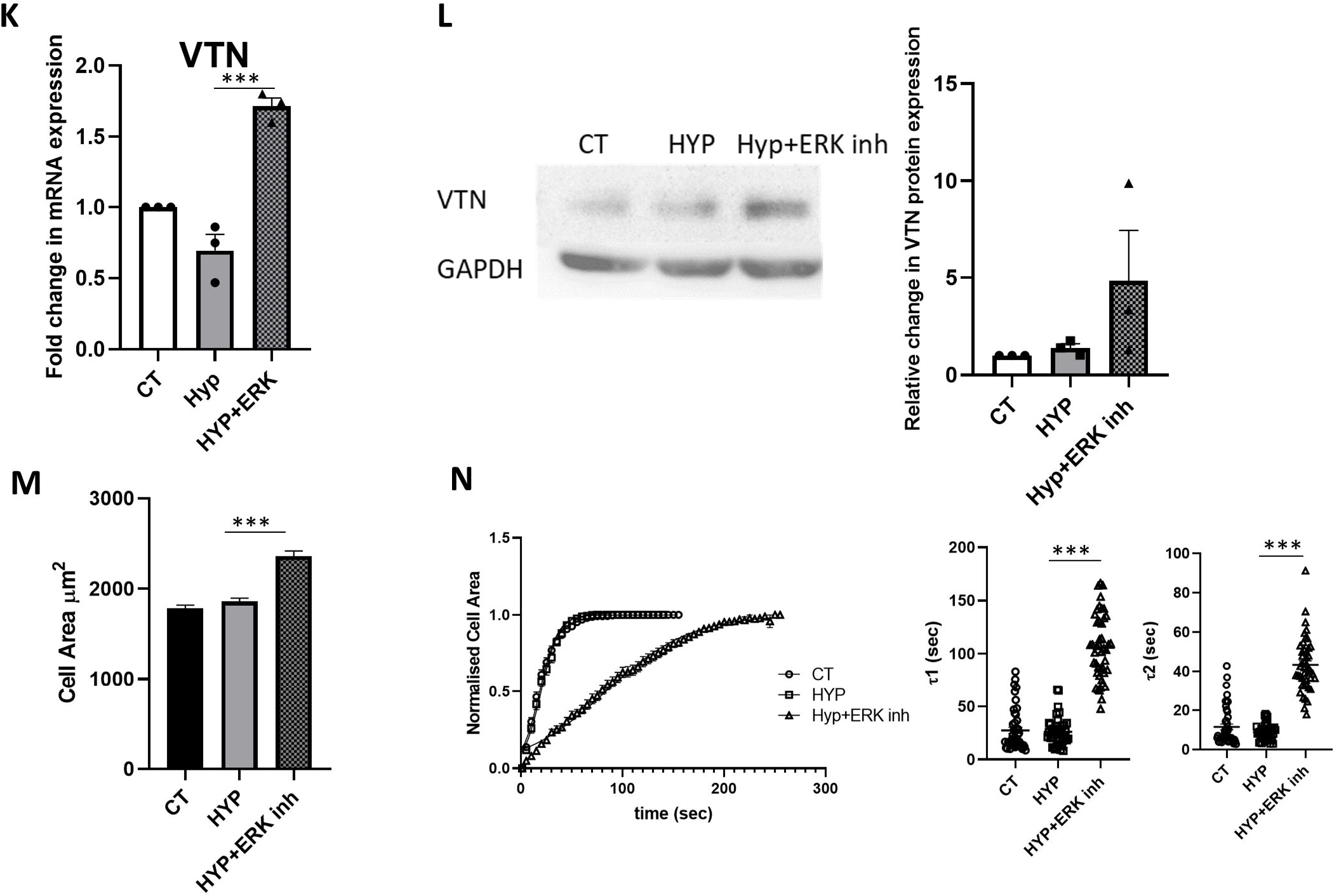
Effect of hypoxia on morphology, adhesion and migration of WJ-MSCs and its co-relation with VTN expression. Cell morphology and cell spread area were compared between WJ-MSCs cultured under control and hypoxia (2% O_2_) conditions. Scale: 500µm (A, B). A representative Gaussian distribution plot reflecting cell spread area comparison for 50 cells is shown (n=3) (C). Cellular de-adhesion dynamics under hypoxia was studied via time lapse imaging of hypoxic WJ-MSCs following EDTA treatment. The normalized cell area vs time data was fitted in Boltzmann sigmoidal equation to obtain the τ1 and τ2 time constant values for a total of 45 cells from three independent biological samples (D). To analyse the cellular migration of WJ-MSCs under hypoxia condition, migration rate (E), cell trajectories (F) and directionality ratio (G) were quantified from DIC time lapse imaging of a total of 60 cells from two independent biological samples. mRNA (H) and protein expression levels (I) of VTN were examined under hypoxia condition and compared against control condition from three independent biological samples. GAPDH was considered as an internal control for both mRNA and protein quantifications (n=3). Actin cytoskeleton profile of control and hypoxia treated WJ-MSCs were visualized via immunofluorescence using Alexa Fluor 488-phalloidin (green) while DAPI (blue) was used to stain nuclei. Representative immunofluorescence images (at 40X magnification) and immunofluorescence mean intensity quantification plot demonstrated no significant change in actin cytoskeleton profile (n=3). Scale: 50µm (J). Effect of ERK pathway inhibition with FR180204 on WJ-MSCs under hypoxia condition. WJ-MSCs were treated with FR180204 for 48h under hypoxia condition. The mRNA and protein expression of VTN was quantified by qRT-PCR and Western blotting, respectively (K, L), in WJ-MSCs treated with or without ERK inhibitor, FR180204. The change in cellular morphology was quantified in terms of cell spread area and plotted from 150 cells obtained from 3 independent biological samples (M). De-adhesion dynamics of WJ-MSCs under hypoxia condition, with or without ERK inhibitor, FR180204 was studied via time lapse imaging. The normalized cell area was fitted in Boltzmann sigmoidal equation and the time constants τ1 and τ2 were plotted for a total of 45 cells from 3 independent biological samples (N). Each bar represents mean ± SEM. (\**p*< 0.05, \*\**p*< 0.01, \*\*\**p*< 0.001).

Interestingly, inhibiting the ERK pathway (negative regulator of VTN) with small molecule inhibitor FR180204, led to induction in VTN expression both at mRNA and protein levels in WJ-MSCs even under hypoxia condition (Fig. 6K, L, *p <* 0.001), and corresponding to the increase in VTN expression, the cells attained flattened morphology with increase in cell spread area (Fig. 6M, *p <* 0.001) and cellular adhesion (Fig. 6N, *p <* 0.001). The significant increase in τ1 and τ2 values with ERK pathway inhibition under hypoxia condition also established the increase in cellular adhesion (Fig. 6N, *p <* 0.001).

Additionally, it was also noted that treating hypoxic WJ-MSCs with VTN enriched conditioned medium collected from ERK pathway inhibited serum starved WJ-MSCs resulted in increased cell spread area (Supp. Fig. 1C, *p <* 0.05) and cellular adhesion (Supp. Fig. 1D, *p <* 0.001) as compared to untreated hypoxic cells. Also, there was a corresponding induction in *VTN* mRNA expression in WJ-MSCs on exposure to this medium as compared to cells treated with VTN knocked down conditioned medium (Supp. Fig. 1E, *p <* 0.05). Altogether, these data implied that the upregulation in VTN expression in WJ-MSCs possibly involved an autocrine signalling mechanism, and that the presence of VTN was responsible for the alteration in adhesion and migration observed in WJ-MSCs under certain stress conditions.

## Discussion

Post transplantation, survival, efficient migration and engraftment of the MSCs in adequate numbers are the key to effective MSC-based therapy. However, exposure to hostile microenvironment prevalent in the damaged or diseased tissue, hamper these features [26]. Hence, exploring the key factors and underlying mechanisms involved in migration of MSCs under hostile micro-environment becomes critical.

Nutrient starvation is one of the major micro-environment stress conditions involved with injury and inflammation in vivo, which could affect retention and functionality of MSCs. In MSCs, serum deprivation has been reported to reduce cellular proliferation and upregulate gene expression associated with maintenance of stemness, angiogenesis and endothelial differentiation [27]. A previous publication from our group showed that when exposed to serum starvation stress for up to 48 hours, WJ-MSCs maintained their viability comparable to control conditions [17]. A study based on preconditioning of bone marrow MSCs with hypoxia and serum deprivation for treatment of ischemic perfusion injury showed that migration of MSCs got reduced on exposure to such stress conditions [28]. However, another contrasting report had shown that serum-deprived MSCs secreted anti-apoptotic factors and had 3-fold increased cellular motility compared to control MSCs [29].

Our current study with WJ-MSCs established that exposure to serum starvation for 48h led to significant increase in cell spread area along with increase in cellular adhesion. In parallel, serum starvation reduced the migration rate, and adversely affected mean square displacement and directionality ratio of WJ-MSCs. Adhesion and migration of cells are profoundly regulated by the interaction of the cells with their surrounding microenvironment via the ECM signalling [30]. Direct coupling between the cell and ECM at the integrin-based adhesion sites allowed the cells to alter their mechanism of migration based on the changes sensed in the surrounding environment [31]. Previous study of our group showed that altered adhesion and migration of WJ-MSCs under febrile temperature stress corresponded to changes in expression of ECM genes like collagens, MMPs and VTN [18]. Here we have investigated the possible role and mechanism of action of ECM gene-VTN in regulation of adhesion and migration of WJ-MSCs under serum starvation stress. VTN is an adhesive multifunctional glycoprotein, known to promote cell adhesion, migration and matrix degradation by binding to integrins, PAI and uPAR, thereby facilitating tissue repair and regeneration [11]. In serum-starved WJ-MSCs, we observed that expression of VTN got upregulated, and as expected, led to increase in adhesion with a corresponding reduction in migration of WJ-MSCs.

Cell adhesion and migration are a coordinated and regulated phenomenon which gets impacted by extracellular signals activating different signal transduction pathways via various surface receptors. Integrin mediated activation of ERK signalling had been shown to positively regulate migration of MSCs via activation of FAK pathway [32, 33]. A recent report showed that IL-1β-induced migration of human umbilical cord derived MSCs (hUC-MSCs) was attenuated by inhibition of ERK, JNK and Akt pathways [34]. In our study with serum-starved WJ-MSCs, the NF-kβ signalling pathway was found to be a positive regulator of VTN expression. And as expected, inhibition of NF-kβ pathway via the p65 knockdown approach resulted in reduction of cellular adhesion and rescue in impaired migration. In addition, ERK signalling pathway appeared to be a negative regulator of VTN in serum starved WJ-MSCs. Thus, ERK pathway inhibition resulted in further upregulation of VTN expression, which led to further increase in adhesion and reduction in migration, highlighting a significant role of VTN in regulating adhesion and migration of WJ-MSCs under serum starvation.

The correlation between increased VTN expression and increased adhesion along with reduced migration was further validated via VTN knockdown studies. Knockdown of VTN led to similar reversal in adhesion and migration patterns as observed with p65 knockdown studies under serum starvation. Simultaneously, there was reduction in mRNA expression of collagens, known contributors of ECM stiffness, which were upregulated under serum starvation treatment alone. A study based on migration of smooth muscle cells on different substrates demonstrated that random migration depended on ECM stiffness in a biphasic manner, whereby beyond an optimal stiffness cell migration got reduced [35]. Thus, the observed alteration in migration pattern in WJ-MSCs on serum starvation treatment might be due to the remodelling of ECM via VTN activated signalling cascade leading to increase in stiffness and cellular adhesion.

Knockdown of VTN also led to the reduction in the number of classical (2-6µm) and super mature (> 6μm length) FAs associated with extensive actin stress fibres formed on exposure to serum starvation. Earlier literature showed that lowering the serum levels in culture medium led to induction in alpha-smooth muscle actin expression and induction of thicker stress fibre formation associated with super mature FAs. The study also showed that increase in super mature FAs led to increased cellular adhesion and reduced turnover of FA in the myofibroblasts [24]. Hence, in our study increase in VTN expression under serum starvation stress appeared to have mediated prominent stress fibre formation along with increase in number of super mature FA.s. This possibly affected the FA turnover due to stronger adhesion of WJ-MSCs which in turn impaired the migration capacity of the cells. This observation was further supported by our ERK pathway inhibition study under serum starvation, wherein corresponding to further increase in VTN expression and stronger reduction in cell migration, formation of extensive stress fibre with super mature FAs were observed to be significantly greater.

Another previous report had shown that FA size could uniquely predict cell speed and these two parameters had a tight bi-phasic relationship, whereby, increase in FA size beyond a threshold resulted in reduction of cell speed as observed in human fibrosarcoma cells [36]. Hence, along with FA length, we quantified FA area which also led to similar observation. Under serum starvation, there was an increase in the number of larger area FAs which got reduced with VTN knockdown, while ERK inhibition under serum starvation increased the number further. Overall, based on our results, VTN appeared to be an important ECM player which could modulate migration of WJ-MSCs under stress condition like serum starvation by altering the cytoskeleton arrangement and FA content and distribution.

To further affirm the role of VTN as the master regulator of the altered physiological attributes, the adhesion and migration pattern of WJ MSCs were evaluated under hypoxia stress, a natural micro-environment condition associated with stem cell niche. Previous reports had shown that hypoxia treatment improved physiological behaviour of MSCs supporting their growth and migration, which was in contrast to the impact of serum starvation [37]. We observed almost no induction in VTN gene expression in WJ-MSCs exposed to hypoxia (2% O_2_). Moreover, WJ-MSCs exposed to hypoxia condition had similar actin cytoskeleton arrangement along with adhesion and migration pattern as control WJ-MSCs. Interestingly, induction of VTN under hypoxia by ERK pathway inhibition led to increase in both cell spread area and adhesion.

To conclude, throughout our study it was noted that induction of VTN under serum starvation stress led to increase in cell spread area and adhesion in WJ-MSCs, while cell migration got reduced. VTN mediated signalling possibly altered the actin cytoskeleton, focal adhesion formation and ECM gene expression which affected the downstream adhesion and migration properties of WJ-MSCs (Fig. 7). Our study, thus, established VTN as the key factor involved in regulating adhesion and migration of WJ-MSCs under certain physiological stress conditions.

**Figure 7:**
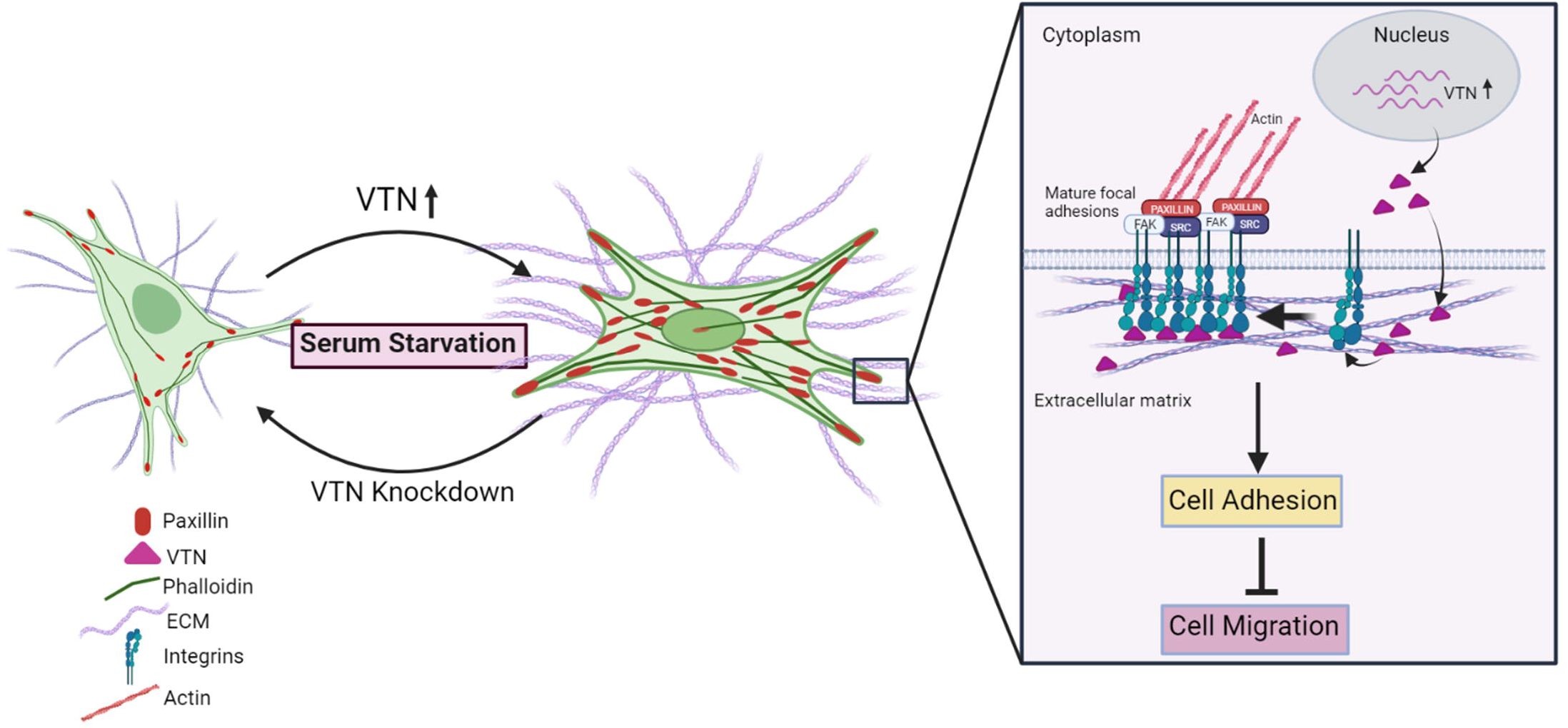
A model representing the impact of serum starvation on adhesion and migration characteristics of WJ-MSCs. Serum starvation stress results in impaired cellular migration of WJ-MSCs via increased cellular adhesion with vitronectin as a key molecular player.

## Acknowledgement

The work was funded by SERB, DST India and IISER, Kolkata. We thank UGC, India for the fellowship of Ms Ankita Sen. We are grateful to Dr. Jayanta Chatterjee, Astha, Kalyani for generously providing umbilical cord samples and Dr. S.S. Jana of IACS, Kolkata for providing the anti-paxillin antibody. We thank Mr. Ritabrata Ghosh for technical assistance with microscopy. We thank Ms Arya Sachan, Mr. Umesh Goyal, Ms Srishti Dutta Gupta for their help with cell culture.

