## Supplemental Fig 1 for "Vitronectin acts as a key regulator of adhesion and migration in human umbilical cord-derived MSCs under different stress conditions"

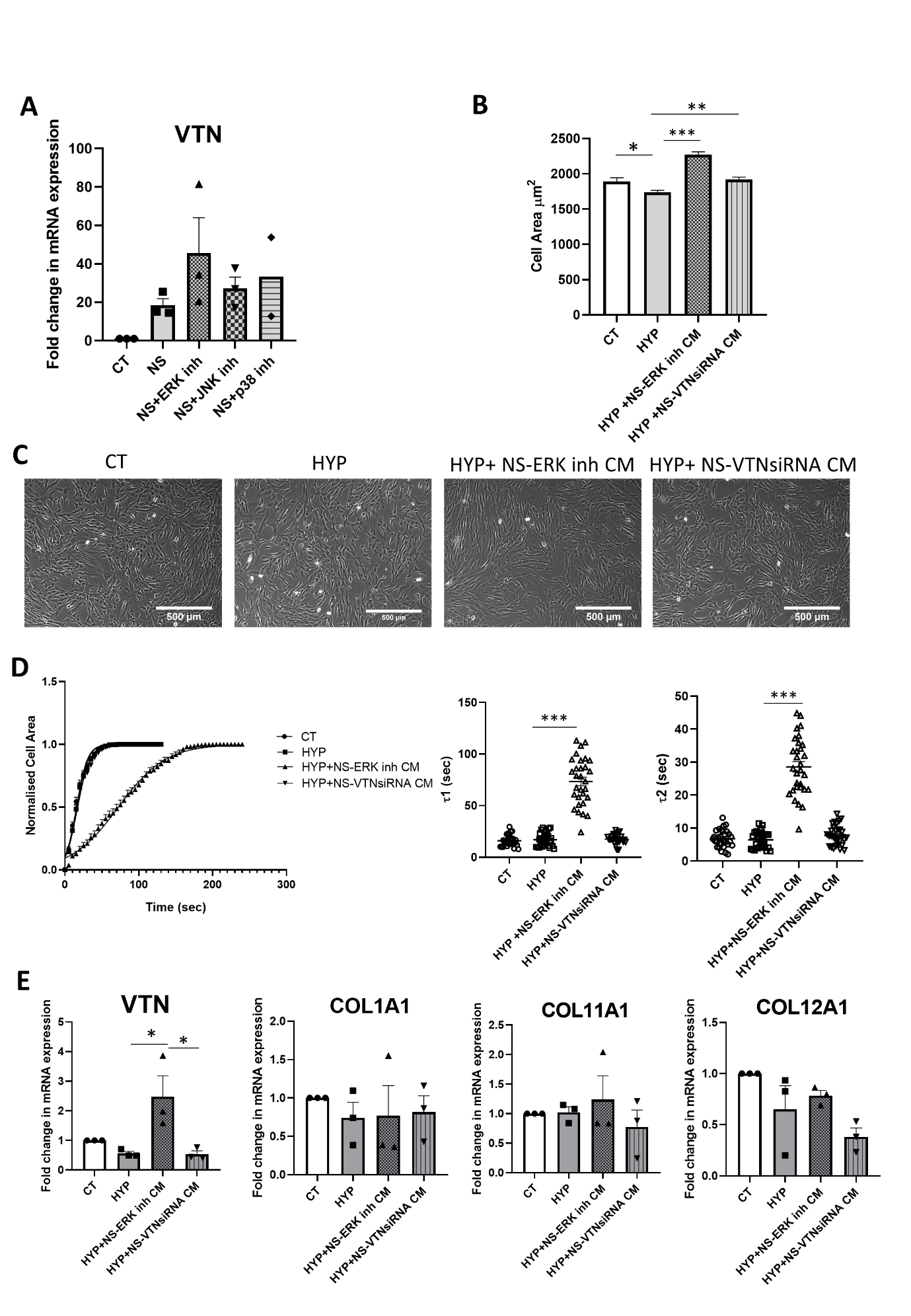

Supplementary figure 1: mRNA expression of VTN, under serum starvation with different signalling pathway inhibition, was detected using qRT-PCR (n=3) (A). Treatment of hypoxic WJ-MSCs with VTN enriched conditioned medium collected from ERK pathway inhibited serum starved WJ-MSCs displayed increase in cell spread area compared to untreated hypoxic cells or VTN knocked down serum starved condition medium treated cells. Bar plot represents comparison of cell spread area from n=100 cells obtained from two independent biological samples (B). Representative morphology images of WJ-hypoxic MSCs treated with conditioned medium collected from ERK inhibited serum starved WJ-MSCs and VTN knocked down serum starved WJ-MSCs. Scale: 500µm (C). The de-adhesion dynamics data of hypoxic WJ-MSCs with the above-mentioned conditioned media treatment was fitted in Boltzmann sigmoidal equation and time constants τ1 and τ2 were plotted for a total of 30 cells from 2 independent biological samples (D). mRNA expression of *VTN* and ECM genes like *COL1A1, COL11A1 and COL12A1* was quantified by qRT-PCR in hypoxic WJ-MSCs with the above-mentioned conditioned media treatment (n=3) (E). Each bar represents mean ± SEM (**p*< 0.05, ***p*< 0.01, ****p*< 0.001).

Supplementary Table 1: Primer sequences used for quantitative reverse transcriptase polymerase chain reaction:

| Gene | Forward (5’-3’) | Reverse (5’-3’) | Amplicon(bp) |
| --- | --- | --- | --- |
| Col1A1 | GGACACAATGGATTGCAAGG | TAACCACTGCTCCACTCTGG | 460 |
| Col11A1 | AGCACAGACGGAGGCAAAC | AGAAGATCAGAATCCCTGCCG | 234 |
| Col12A1 | ACCCACCTTCCGACTTGAATT | TAGGCCCATCTGTTGTAGGG | 122 |
| GAPDH | GAGTCAACGGATTTGGTCGT | TTGATTTTGGAGGGATCTCG | 248 |
| MMP-1 | CCAGTATGCACAGCTTTCCTC | TGCCTCCCATCATTCTTCAGG | 154 |
| VTN | GGGTCTACTTCTTCAAGGGGAA | AATGAACTGGGGCTGTCTGG | 197 |

Supplementary Video 1: DIC live cell imaging of WJ-MSCs under control condition (Fig. 1E).

Supplementary Video 2: DIC live cell imaging of WJ-MSCs under serum starvation condition (Fig. 1E).

Supplementary Video 3: DIC live cell imaging of WJ-MSCs under control condition (Fig. 4E).

Supplementary Video 4: DIC live cell imaging of WJ-MSCs with NC siRNA treatment under serum starvation condition (Fig. 4E).

Supplementary Video 5: DIC live cell imaging of WJ-MSCs with VTN siRNA treatment under serum starvation condition (Fig. 4E).
